# Childhood conduct problems are associated with reduced white matter fibre density and morphology

**DOI:** 10.1101/2020.05.29.123364

**Authors:** Daniel T. Burley, Sila Genc, Timothy J. Silk

## Abstract

Childhood conduct problems are an important public health issue as these children are at-risk of adverse outcomes. Studies using diffusion Magnetic Resonance Imaging (dMRI) have found that conduct problems in adults are characterised by abnormal white-matter microstructure within a range of white matter pathways underpinning socio-emotional processing, while evidence within children and adolescents has been less conclusive based on non-specific diffusion tensor imaging metrics. Fixel-based analysis (FBA) provides measures of fibre density and morphology that are more sensitive to developmental changes in white matter microstructure. The current study used FBA to investigate whether childhood conduct problems were related both cross-sectionally and longitudinally to microstructural alterations within the fornix, inferior fronto-occipital fasciculus (IFOF), inferior longitudinal fasciculus (ILF), superior longitudinal fasciculus (SLF), and the uncinate fasciculus (UF). dMRI data was obtained for 130 children across two time-points in a community sample with high levels of externalising difficulties (age: time-point 1 = 9.47 – 11.86 years, time-point 2 = 10.67 −13.45 years). Conduct problems were indexed at each time-point using the Conduct Problems subscale of the parent-informant Strengths and Difficulties Questionnaire (SDQ). Conduct problems were related to lower fibre density in the fornix at both time-points, and in the ILF at time-point 2. We also observed lower fibre cross-section in the UF at time-point 1. The change in conduct problems did not predict longitudinal changes in white-matter microstructure across time-points. The current study suggests that childhood conduct problems are related to reduced fibre-specific microstructure within white matter fibre pathways implicated in socio-emotional functioning.

## Introduction

Antisocial behaviour is a behavioural phenotype that includes violence and aggression, difficulties following rules, and behaviours that violate the rights of others. Antisociality is associated with a range of negative outcomes including crime, imprisonment, psychopathology, substance misuse, lower educational attainment and poorer physical health (Fergusson, John Horwood, & Ridder, 2005; Moffitt & Scott, 2008; Mordre, Groholt, Kjelsberg, Sandstad, & Myhre, 2011; Odgers et al., 2007; Odgers et al., 2008). There are also a number of adverse impacts of antisocial behaviour on their victims, as well as wider negative economic and societal costs, meaning that antisocial behaviour is viewed as a public health issue (National Institute for Health and Care Excellence (NICE), 2013). Importantly, antisociality is increasingly understood as a multifaceted neurodevelopmental construct with conduct problems emerging in childhood and adolescence (Fairchild, Van Goozen, Calder, & Goodyer, 2013; Piquero, Farrington, Nagin, & Moffitt, 2010; Raine, 2018).

There is considerable evidence that antisociality is characterised by emotional processing impairments, such as reduced empathy, lower physiological affective responsivity, diminished capacity to learn about punishment and reward, and emotional dysregulation (K. Blair, Morton, Leonard, & Blair, 2006; R. Blair, 1999; Fanti et al., 2019; Gao, Raine, Venables, Dawson, & Mednick, 2010; Hunnikin, Wells, Ash, & Van Goozen, 2019; Van Goozen, Fairchild, Snoek, & Harold, 2007; Van Langen, Wissink, Van Vugt, Van der Stouwe, & Stams, 2014). Using magnetic resonance imaging (MRI), studies have identified structural and functional abnormalities in limbic brain regions important for processing emotion, in particular the amygdala, and prefrontal regions implicated in affective decision-making, learning and regulation, such as the orbitofrontal and ventromedial prefrontal cortex (Baker, Clanton, Rogers, & De Brito, 2015; Coccaro, McCloskey, Fitzgerald, & Phan, 2007; Meyer-Lindenberg et al., 2006; Noordermeer, Luman, & Oosterlaan, 2016; Pardini, Raine, Erickson, & Loeber, 2014; Rogers & De Brito, 2016; Thomas et al., 2012). In addition, individuals high in antisocial behaviour demonstrated reduced functional connectivity between limbic and prefrontal regions (Contreras-Rodríguez et al., 2015; Finger et al., 2012; Fulwiler, King, & Zhang, 2012; Stoddard et al., 2017). Theories of antisociality have therefore proposed that antisocial behaviour reflects dysfunction in neural networks implicated in emotional processing and learning, including disrupted connections between limbic and prefrontal regions (R. Blair, 2005; Kiehl, 2006; Raine, 2018).

More recent research has used diffusion tensor imaging (DTI) to examine white matter microstructure between brain regions within individuals high in antisociality. DTI measures such as fractional anisotropy (FA) are sensitive to the anisotropic organisation of a white matter fibre, whereas mean diffusivity (MD) can represent the mean isotropic diffusion within the extracellular space. Given the theorised abnormality between limbic and prefrontal regions in relation to antisociality, there has been much focus on examining the microstructure of the uncinate fasciculus (UF) - a long range white matter pathway that connects limbic regions within the temporal lobe to frontal regions. Studies have found that adults with antisocial behaviour have reduced white matter organisation in the UF (Craig et al., 2009; Hoppenbrouwers et al., 2013; Sundram et al., 2012). Beyond the UF, studies have found evidence that adults high in antisociality show diminished white matter organisation in additional association pathways including the inferior fronto-occipital fasciculus (IFOF), inferior longitudinal fasciculus (ILF), superior longitudinal fasciculus (SLF), and fornix (Bolhuis et al., 2019; Hoppenbrouwers et al., 2013; Karlsgodt et al., 2015; Lindner et al., 2016; Sethi et al., 2015; Sundram et al., 2012). This suggests that increased antisociality may be reflected in widespread altered white matter architecture that extend beyond specific impairments in the UF.

In terms of antisociality in childhood and adolescence, studies have explored similar white matter pathways to examine whether these microstructural differences are evident earlier in development. This is important to understand as the brain shows rapid development across adolescence and offers a window of plasticity during which interventions may be able to shape developmental trajectories (Blakemore, 2012; Lebel & Beaulieu, 2011). However, neuroimaging studies investigating conduct problems in children and adolescents have produced mixed findings. While there is evidence for greater white matter microstructural organisation in adolescents with conduct difficulties across association tracts (Breeden, Cardinale, Lozier, VanMeter, & Marsh, 2015; Decety, Yoder, & Lahey, 2015; Haney-Caron, Caprihan, & Stevens, 2014; Li, Mathews, Wang, Dunn, & Kronenberger, 2005), other studies have reported lower microstructural organisation (Grazioplene et al., 2020; Passamonti et al., 2012; Peper, De Reus, Van Den Heuvel, & Schutter, 2015; Sarkar et al., 2013; Zhang, Zhu, et al., 2014) or no difference compared to adolescents without conduct problems (Decety et al., 2015; Finger et al., 2012; Hummer, Wang, Kronenberger, Dunn, & Mathews, 2015; Passamonti et al., 2012; Puzzo et al., 2018; Sarkar et al., 2013; Zhang, Gao, et al., 2014; Zhang, Zhu, et al., 2014).

Many of these studies recruited youths with wide age ranges, which may contribute to inconsistent findings given that white matter microstructure develops in a time-dependent fashion within major white-matter tracts (Lebel & Beaulieu, 2011; Lebel, Walker, Leemans, Phillips, & Beaulieu, 2008). In addition, the effects of cooccurring forms of psychopathology could influence white matter microstructure given the comorbidity of conduct problems with alternative externalising and internalising difficulties (Lahey et al., 2008; Patalay et al., 2015), which may have contributed to the inconsistent literature. It is therefore important to examine the effect of conduct problems alongside additional forms of psychopathology to investigate the specificity of altered white matter microstructure.

One further potential reason for the contrasting results observed in children and adolescents may be due to the metrics previously used in DTI studies to index white matter microstructure. Measures such as FA and MD are relatively non-specific at distinguishing between specific fibre properties, such as axon density, crossing fibres and myelination, which are separate physio-anatomical white matter properties important for understanding developmental changes (Beaulieu, 2014). In contrast, fixel-based analysis (FBA) (Raffelt et al., 2012; Raffelt et al., 2017) is a recently developed advanced diffusion MRI analysis technique that uses fibre-specific information to model the intracellular space within white matter and provide specific measures of axon density and fibre bundle cross-section. FBA produces metrics that index fibre density (FD), which represents the intra-axonal volume fraction of white matter fibres, fibre cross-section (FC), which refers to the cross-sectional area of voxels that a fibre occupies, as well as the combined effect of fibre density and cross-section (FDC) (Raffelt et al., 2017). These indices aim to quantify and disentangle fibre-specific white matter properties more accurately compared to more traditional DTI metrics such as FA. Importantly, studies have demonstrated that FBA is more sensitive to developmental changes in white matter architecture (Kelley, Plass, Bender, & Polk, 2019).

Grazioplene et al. (2020) is the only study to date to have used fixel-based analysis fibre to examine childhood conduct problems and white matter microstructure. Using a cross-sectional design, a group of 70 children with parent-rated aggressive behaviour were compared to matched controls aged 8 – 16 years old for mean FD across a range white matter tracts. Children showing aggression demonstrated lower FD in a cluster of limbic and cortical pathways including the IFOF and fornix relative to controls, increased FD in the corpus callosum, and dimensional analysis revealed an association between aggression and reduced FD in the cingulum bundle. The current study intended to extend this study and previous research to investigate childhood conduct problems in relation to fibre density *and* morphology as both measures are sensitive to changes during development (Genc et al., 2018), and to examine these relationships within a longitudinal design that allowed us to also examine the effects of conduct problems on white matter microstructural development.

### Current Study

We investigated conduct problems and white matter microstructure in a large community-based sample of children aged 9-13 across two time-points specifying narrow age ranges. We implemented FBA to investigate fibre density and morphology within the fornix, IFOF, ILF, SLF and UF, which are all implicated within socio-emotional processing systems (Ameis & Catani, 2015) and have been linked with antisociality (Waller, Dotterer, Murray, Maxwell, & Hyde, 2017). Importantly, the age-ranges were narrow at each time-point across participants to examine white-matter microstructure at specific developmental stages and to allow us to precisely explore developmental changes across time. We also considered cooccuring dimensions of psychopathology to examine the specificity of white matter microstructure effect to conduct problems. We hypothesised that conduct problems would be cross-sectionally associated with lower FD, with no relationship observed for FC, consistent with research that has linked neurodevelopmental difficulties with decreased white matter fibre density in adolescence, rather than lower macroscopic cross-section of fibres (Dimond et al., 2019; Genc et al., 2020). We also predicted longitudinal relationships such that change in conduct problems across time-points would be associated with FD development within each tract, with no relationship emerging for FC development.

## Method

### Participants

Participants were recruited as part of the Neuroimaging of the Children’s Attention Project (NICAP; Silk et al. 2016), an Australian longitudinal multimodal neuroimaging study of community-based cohort of children with and without Attention Deficit Hyperactivity Disorder (ADHD). This longitudinal study was approved by The Royal Children’s Hospital Melbourne Human Research Ethics Committee (HREC #34071). Written informed consent was obtained from the parent/guardian of all children enrolled in the study. Children were excluded from the study if they had a neurological disorder, intellectual disability, or serious medical condition (e.g. diabetes, kidney disease).

Full details of the NICAP cohort and assessment methods are detailed in Silk et al. (2016). Briefly, children were initially recruited from 43 socio-economically diverse primary schools distributed across the Melbourne metropolitan area, Victoria, Australia (Sciberras 2013), and underwent comprehensive assessment for ADHD at age 7 via the Diagnostic Interview Schedule for Children (DISC-IV) completed with parents face-to-face (Sciberras et al. 2013). Children were categorised as either meeting a negative or positive diagnosis for ADHD. At a 36-month follow-up, a subset of participants were invited for an appointment at The Melbourne Children’s campus, which included a child assessment, parent questionnaire, mock scan, and MRI scan at age 10 (subsequently referred to as time-point 1). Youths with ADHD represent an at-risk adolescent sample for conduct problems given the high comorbidity between ADHD and additional externalising difficulties including oppositional defiant disorder (ODD) and conduct disorder (CD) (Beauchaine, Hinshaw, & Pang, 2010; R. Blair, White, Meffert, & Hwang, 2013; Lahey et al., 2008). The DISC-IV was repeated at time-point 1 to re-assess ADHD group status, as well as to examine ODD status (see Table 1); 40% of the children met diagnostic criteria for ADHD, 30 % met criteria for ODD, and 6.9 % met criteria for CD. Children were invited for a follow-up appointment (subsequently referred to as time-point 2) approximately 16 months following their initial visit (M = 16.08, SD = 2.32 months). Overall, only data from the two imaging time-points: time-point 1 (age: M = 10.38, SD = 0.44 years old) and time-point 2 (age: M = 11.72, SD = 0.51 years old); were included for analysis in the current study. Direct assessments and MRI scans were performed by a trained research assistant blind to the child’s diagnostic status.

**Table 1.**
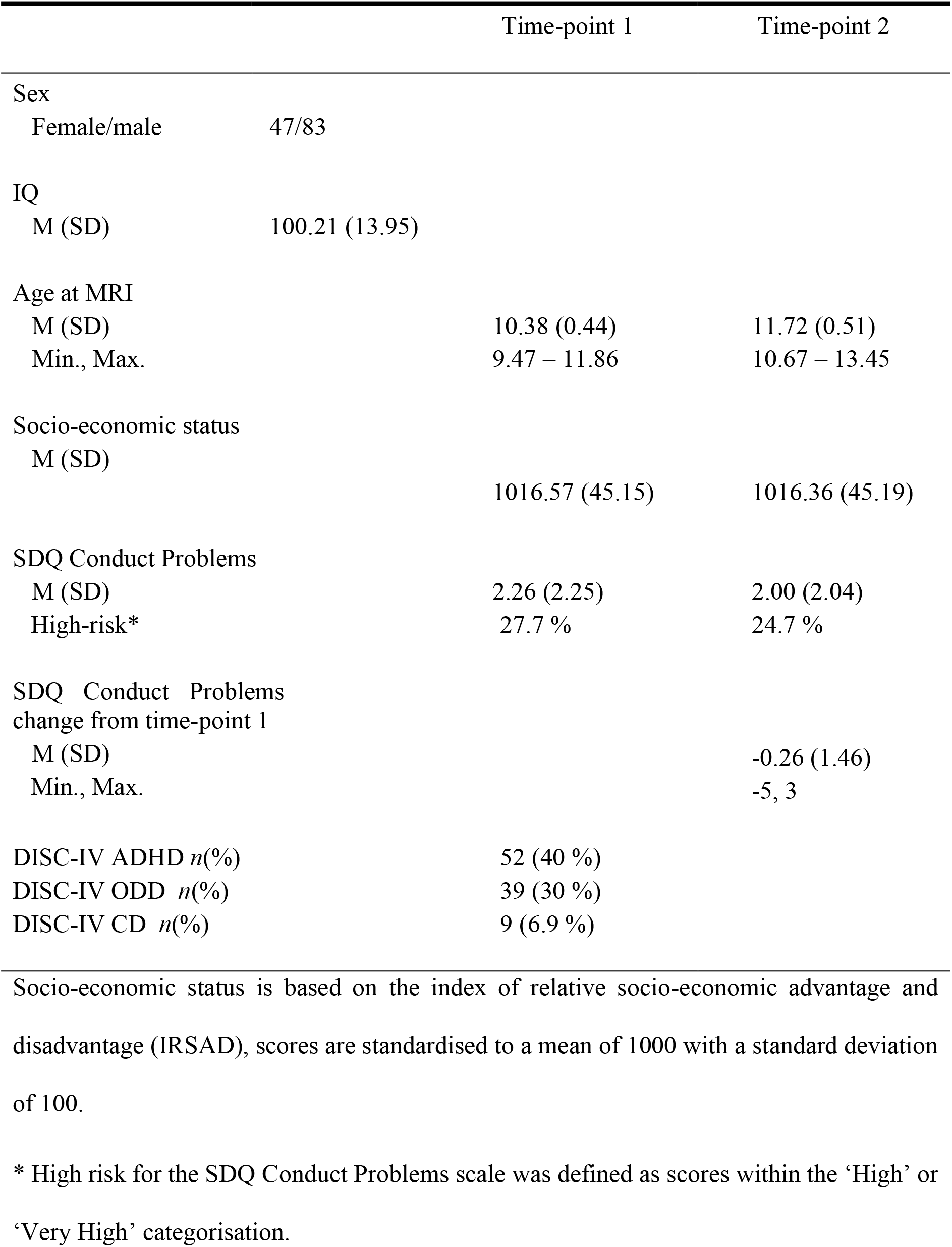
Demographic information for the sample across imaging time-points.

Participant’s socio-economic status was indexed based on scores from the Index of Relative Socio-economic Advantage and Disadvantage (IRSAD) taken from the Socio-Economic Indexes for Areas obtained at each time-point from the Australian Bureau of Statistics (https://www.abs.gov.au/websitedbs/censushome.nsf/home/seifa). IRSAD scores were developed based on several variables for a given area, including income, education, unemployment, and are standardised to a mean of 1000 with a standard deviation of 100. Participant’s estimated full-scale intellectual functioning (IQ) was assessed at a previous assessment at age 7 (Mage = 7.23, SDage = 0.38) using the vocabulary and matrix reasoning subtests of the Wechsler Abbreviated Scale of Intelligence (WASI (Wechsler, 1999).

### Conduct problems

Conduct problems were indexed using the Conduct scale of the parent-rated Strengths and Difficulties Questionnaire (SDQ; Goodman, 1997) completed at time-point 1 and 2. The SDQ is a 25-item screening tool that assess children across several areas of functioning including conduct, emotional, hyperactivity/inattention, interpersonal problems, as well as examining prosocial behaviours. The conduct problems scale includes 5-items (e.g. “Often fights with other children”, “Often lies or cheats” and “Steals from home, school or elsewhere”) scored across 0 (Not true), 1 (Somewhat true), and 2 (Certainly true).

### Magnetic resonance imaging (MRI)

Diffusion MRI data were acquired at two distinct time-points on a 3.0 T Siemens Tim Trio, at The Melbourne Children’s Campus, Parkville, Australia. Data were acquired using the following protocol: b = 2800 s/mm^2^ (60 directions), voxel-size = 2.4×2.4×2.4 mm, echo-time / repetition time (TE/TR) = 110/3200 ms. A total of 152 participants had longitudinal MRI data. Of those, 130 participants had useable diffusion MRI data at both time-points, therefore the subsequent image processing and analysis was performed on these 130 participants with imaging data at two time-points. All dMRI data were processed using MRtrix3 (v3.0RC3; Tournier et al., 2019) using pre-processing steps from a recommended longitudinal fixel-based analysis (FBA) pipeline (Raffelt et al., 2017; Genc et al., 2018). Full details of processing and analysis steps are listed in Genc et al. (2020). Briefly, images were denoised, corrected for motion, eddy current and susceptibility induced distortions, bias field corrected, and upsampled. Fibre orientation distributions were computed using a group average response function, and maps were subsequently registered to a common template.

Images were visually inspected for motion artefact (assessed by the presence of Venetian blinding artefact), and whole datasets were excluded if excessive motion was present. In addition, we calculated mean frame-wise displacement using the FSL software library (v5.0.10) (Smith et al. 2004).

### White matter tract dissection

We chose to delineate five bilateral fibre pathways that have been previously found to show diminished white matter organisation for individuals with antisocial behaviour: Inferior fronto-occipital fasciculus (IFOF); superior longitudinal fasciculus (SLF); inferior longitudinal fasciculus (ILF); uncinate fasciculus (UF); and the fornix (FX) in our population template space. First, we transformed and warped tractography masks from the JHU-ICBM atlas (Smith et al., 2004) to our longitudinal template. Then, we placed anatomically informed regions of interest (ROIs) from a defined protocol (Wakana et al., 2007) ensuring that these regions overlapped with both the warped tractography masks as well as the whole brain tractogram. Finally, we segmented fixels (fibre-specific voxels) from the whole-brain template which corresponded with our tracts of interest.

### Statistical Analysis

Statistical analyses were performed within R (version 3.6.2) and visualisations were carried out in RStudio (version 1.2.1335). We tested the cross-sectional relationship between SDQ conduct problems and each FBA metric (mean FD and mean FC) at each time-point. We also assessed the change in conduct problems scores and the change in each FBA metric (termed FD_diff_/FC_diff_) across time-points to assess the longitudinal relationship between conduct problems and white-matter development. For both cross-sectional and longitudinal analyses, linear mixed-effects models were computed using *lme4* for each FBA metric with conduct problems entered as random effects and white-matter tract as fixed effects to explore the main effect of SDQ conduct problems and the interaction between conduct problems and white-matter tract. We also included participant age, IQ, socio-economic status, sex, ADHD diagnostic status, participant in-scanner motion and total intracranial volume (for FC and FC_diff_) as covariates within each model. Examples of the models used for the cross-sectional (1) and longitudinal analysis (2) for mean FD are detailed below:

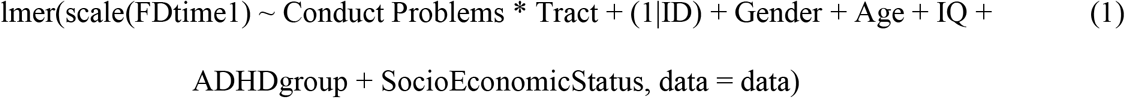

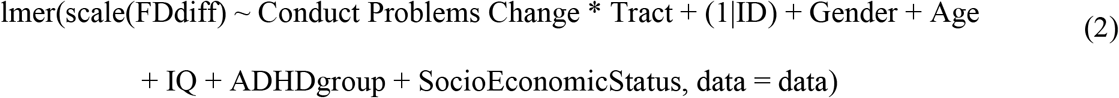

Significant main effects and interactions for conduct scores and white-matter tracts were explored further by running individual correlations between conduct problems scores and the relevant FBA metric for each white matter tract. Within these individual correlations, we covaried only for variables that were predictive in the previous mixed effects model to maintain statistical power. As multiple correlations were run for each FBA metric, *p*-values were adjusted when running the correlational analysis using False Discovery Rate (FDR) correction (Benjamini & Hochberg, 1995). All variables were centred prior to analysis.

## Results

Table 1 details the demographic characteristics of the adolescent sample across time-points. The sample included showed a range of conduct problem scores including 27.7% of children who were rated as high or very high risk of conduct problems at time-point 1 and 24.7% at time-point. Figure 1A and 1B illustrate the age and conduct problems scores for the sample at each time-point. Mean FBA metrics for each white-matter tract are included across time-points in Table 2. Each FBA metric correlated highly across time-points for each tract (*p*s < .001).

**Figure 1.**
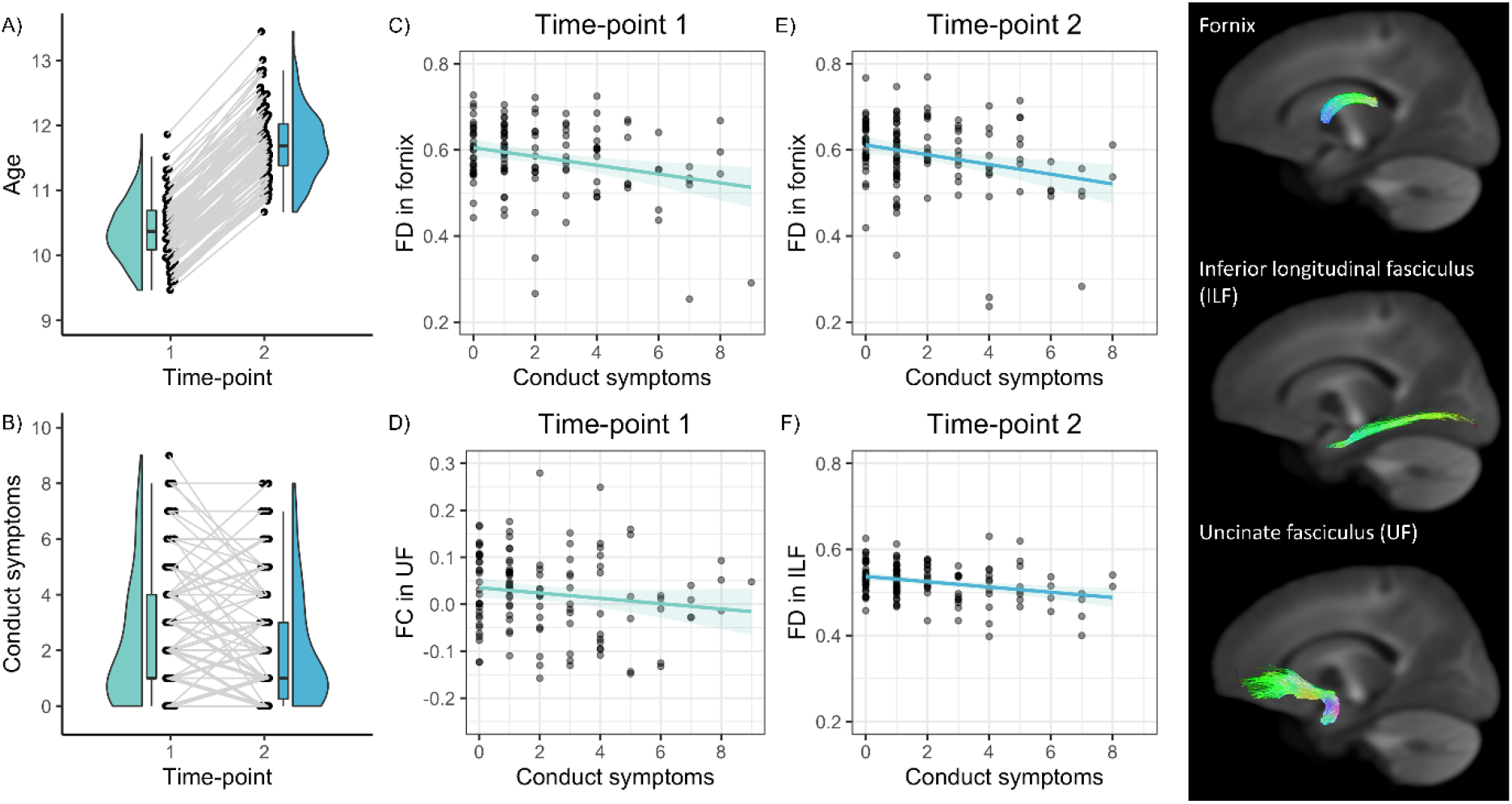
Panels A-B. Participant age and change in SDQ conduct problems over the two time-points. Longitudinal data points are connected by a line. **Figure 1. Panels C-F**. Significant relationships between SDQ Conduct Problems and mean FBA metrics across tracts and time-points: Panel C, fibre density within the fornix at time-point 1; Panel D, fibre density within the inferior longitudinal fasciculus, ILF, at time -point 1; Panel E, fibre density within the fornix at time -point 2; Panel F, fibre density within the uncinate fasciculus at time -point 1. Trend-line is illustrated with 95% confidence interval.

**Table 2:** Descriptive data for fibre density and fibre cross-section for each white-matter tract at each time-points.

|  | Fibre density |  |  | Fibre cross-section |  |  |
| --- | --- | --- | --- | --- | --- | --- |
|  | Time-point 1 | Time-point 2 | Change | Time-point 1 | Time-point 2 | Change |
|  | M (SD) Min, Max | M (SD) Min, Max | M (SD) Min, Max | M (SD), Range | M (SD), Range | M (SD) Min, Max |
| Fx | .58 (.08) .25, .73 | .59 (.09) .24, .77 | .007 (.03) -.08, .13 | -.07 (.10) -.26, .23 | -.06 (.11) -.26, .34 | .011 (.03) -.06, .10 |
| IFOF | .56 (.03) .48, .64 | .57 (.03) .49, .64 | .006 (.01) -.03, .04 | .04 (.07) -.13, .21 | .05 (.07) -.12, .22 | .010 (.02) -.06, .07 |
| ILF | .52 (.05) .38, .65 | .53 (.04) .40, .63 | .006 (.02) -.03, .07 | .08 (.10) -.11, .32 | .12 (.10) -.09, .43 | .015 (.04) -.08, .14 |
| SLF | .54 (.04) .45, .61 | .54 (.04) .47, .63 | .009 (.01) -.02, .04 | .08 (.11) -.19, .32 | .11 (.11) -.15, .35 | .032 (.03) -.08, .18 |
| UF | .56 (.04) .41, .68 | .57 (.04) .44, .70 | .004 (.03) -.07, .07 | .02 (.09) -.28, .28 | .03 (.09) -.24, .28 | .009 (.03) -.10, .12 |

Time-point 1, $n = 125$ ; time-point 2, $N = 130$
Fx, Fornix, IFOF, inferior fronto-occipital fasciculus; ILF, inferior longitudinal fasciculus; SLF, superior longitudinal fasciculus; UF, uncinate fasciculus.

### Cross-sectional analysis

Linear mixed-effects models revealed that there was a main effect of conduct problems for mean FD at time-point 2, *F*(1,122) = 4.74, *p* = .03, but not for the remaining FBA metrics/time-points (*p*s > .10). There was a significant interaction between conduct problems and tract region for mean FD at both time-point 1, *F*(4, 492) = 5.05, *p* < .001, and time-point 2, *F*(4, 512) = 5.77, *p* < .001, as well as for mean FC at time-point 1, *F*(4, 492) = 2.72, *p* = .03. This supported the investigation of tract-specific relationships with respect to conduct problems. There was no interaction between conduct problems and tract region for mean FC at time-point 2, *F*(4, 512) = 1.74, *p* = .14.

To follow up significant interactions, we ran zero-order/partial correlations between conduct problems and FD at both time-points and FC at time-point 1 for each white matter tract individually (see Table 3). We included variables that were significant predictors in the original linear mixed effects models as covariates (total intracranial volume for FC at time-point 1, IQ for FD at time-point 2), although the results were unaltered when controlling for IQ and so the results are reported with this variable removed to maintain statistical power. Conduct problems were related with lower mean FD in the fornix at time-point 1 and 2, as well as the ILF at time-point 2 (*p*_FDR_ < .05). Conduct problems were associated with low mean FC in the UF at time-point 2, although this failed to survive FDR-adjusted statistical significance. Figure 1C-1F illustrates these significant relationships.

**Table 3.** Zero-order and partial correlations between parent-rated SDQ conduct problems and fixel-based analysis metrics across white-matter tracts.

|  | Fibre density |  |  |  | Fibre cross-section |  |
| --- | --- | --- | --- | --- | --- | --- |
|  | Time-point 1 |  | Time-point 2 |  | Time-point 1 |  |
|  | *r* | *p* | *r* | *p* | *r* | *p* |
| Fx | **-.27\*** | **.003** | **-.27\*** | **.002** | -.03 | .72 |
| IFOF | -.13 | .16 | -.17 | .05 | -.06 | .51 |
| ILF | -.14 | .13 | **-.29\*** | **.001** | -.02 | .88 |
| SLF | -.02 | .86 | .01 | .92 | .14 | .12 |
| UF | -.12 | .17 | -.09 | .30 | **-.20** | **.03** |

*Note.* Correlations were run only for metrics and time-points where a significant SDQ conduct problems and white matter tract interaction had been identified within the previous mixed-effects model analysis.
Significant relationships highlighted in bold and associations that survived adjusted statistical significance at $p_{\text{FDR}} < .05$ are annotated with an asterisk (\*)
Time-point 1, $n = 125$ ; time-point 2, $N = 130$
Total intracranial volume entered as a covariate for FC at time-point 1
Fx, Fornix, IFOF, inferior fronto-occipital fasciculus; ILF, inferior longitudinal fasciculus; SLF, superior longitudinal fasciculus; UF, uncinate fasciculus.

### Longitudinal analysis

Linear mixed-effects models revealed that there was no main effect of the change in conduct problems across time-points nor an interaction with tract region for either FD_diff_ and FC_diff_ (*ps* > .22) so no further analyses were conducted for these longitudinal FBA metrics.

### Specificity of cross-sectional findings to childhood conduct problems

Given the comorbidity of neurodevelopmental difficulties including conduct problems (Lahey et al., 2008; Patalay et al., 2015), we also assessed whether conduct problems were driving the observed findings in relation to the SDQ subscales. We ran multiple linear regressions entering all SDQ subscales (conduct, emotional, hyperactivity/inattention, interpersonal problems, and prosocial behaviours) to predict the specific FBA metrics at the relevant time-point that we had previously identified as associated with conduct difficulties. We continued to control for total intracranial volume when predicting FC. The analysis showed that when all SDQ subscales were entered as predictor variables, conduct problems was the primary unique predictor of reduced mean FD for the fornix across both time-points (time-point 1, *t*(5, 119) = −1.95, *p* = .05, *β* = −.24; time-point 2, *t*(5, 124) = −2.28, *p* = .02, *β* = −.29) and within the ILF at time-point 2 (although this did not surpass statistical significance), *t*(5, 124) = −1.83, *p* = .07, *β* = −.23, and the remaining SDQ subscales did not uniquely predict any of these FBA metrics. SDQ Conduct Problems were however not uniquely predictive of mean FC within the UF *t*(6, 118) = −1.35, *p* = .18, *β* = −.13, likewise to the remaining SDQ subscales.

## General Discussion

In this longitudinal study of children aged 9-13 years, we used fixel-based analyses to determine whether conduct problems were related to specific microstructural alterations within several white matter tracts both cross-sectionally and longitudinally. We identified that greater conduct problems were related to lower fibre density in the fornix at both time-points, as well as in the ILF at the second time-point. Longitudinally, the change in conduct difficulties did not predict the development of either fibre density or morphology across time-points. These results partially support our hypotheses as abnormalities related to conduct difficulties were specific to FD, rather than FC, although these alterations were not universal across all white matter tracts and did not extend to altered longitudinal development in FD.

We used advanced analysis techniques to examine specific fibre properties within white matter tracts and identified that conduct problems in childhood were related to reduced fibre density within several association white-matter pathways, which was consistent with Grazioplene et al. (2020). Specifically, we found that conduct difficulties in late childhood/early adolescence were characterised by a lower intra-axonal volume fraction for the fornix and ILF that indicates reduced density of axons along these pathways. The current research extended Grazioplene et al. (2020) to also examine fibre cross-section and found conduct problems were associated with macroscopic cross-section of fibres within the UF specific to the earlier time-point. Alterations to axonal microstructure, by way of reduced axon count or diameter, could result in deficiencies for processing speed and conduction velocity across the brain (Drakesmith et al., 2019; Horowitz et al., 2015), and altered white matter microstructure in these pathways may contribute to the risk of conduct difficulties. We however note that only reduced fibre density within the fornix and ILF were specifically associated with conduct problems when co-occurring dimensions of psychopathology were controlled for, which may indicate that reduced axonal density is the driving property underlying microstructural alterations specific to conduct problems. The current study also examined the longitudinal effect of conduct problems on FBA metrics, but no associations emerged for white matter development. The current study increases our understanding of the specific underlying microstructural properties associated with conduct problems during development.

The fornix, ILF and UF have each been implicated in wider socio-emotional networks (Ameis & Catani, 2015) and therefore developmental microstructural alterations – and potentially inefficient processing - within these networks may reflect emerging socio-emotional impairments observed in relation to conduct difficulties. The UF has been suggested as the key connection within a ‘temporo-amygdala-orbitofrontal’ network (Catani, Dell’Acqua, & De Schotten, 2013) that is critical to the regulation of social and emotional behaviour (Von Der Heide, Skipper, Klobusicky, & Olson, 2013). The ILF also demonstrates connections within the temporal lobe with projections at the posterior temporal lobes and occipital lobes (Catani, Jones, Donato, & Ffytche, 2003). By virtue of these connections with temporal regions, including the amygdala (Fox, Iaria, & Barton, 2008), the ILF has been linked to the integration of visual and emotional processes (Catani, Howard, Pajevic, & Jones, 2002) and both the ILF and UF have been implicated in facial expression processing (Coad et al., 2017; Unger, Alm, Collins, O’Leary, & Olson, 2016), which has been demonstrated to be impaired in antisocial populations (Dawel, O’Kearney, McKone, & Palermo, 2012; Marsh & Blair, 2008).

Conversely, no study in adults have linked antisociality to disrupted microstructure in the fornix, although Breeden et al. (2015) has reported that adolescents with high antisocial behaviour showed reduced FA compared to non-antisocial controls, which was linked to callous symptoms. The fornix represents a major output pathway of the amygdala that projects to the mammillary bodies and hypothalamic regions implicated in fear processing (Walker, Toufexis, & Davis, 2003). The fornix also forms part of the Papez circuit, which is an important structure within the limbic system, and involved in the regulation of emotions by higher order frontal areas (Lövblad, Schaller, & Vargas, 2014). In addition, there is evidence that increased white-matter microstructural organisation within the fornix is associated with elevated anxiety (Modi et al., 2013) and, therefore, diminished white-matter microstructural properties within the fornix may reflect diminished anxiety, consistent with patterns of fearlessness associated with conduct problems and callous symptoms in children (Fanti, 2016).

We note that the relationships between conduct difficulties and altered mean FD were specific to the fornix and ILF. There is evidence that the fornix and ILF show an early peak in FA suggesting earlier developmental maturation compared to many of the other major white-matter tracts (Lebel et al., 2012; Lebel, Treit, & Beaulieu, 2017; Slater et al., 2019). Altered white matter microstructural development may therefore be more evident at a younger age within the fornix and ILF, which may account for the specificity of the current findings in relation to conduct problems. In contrast, white matter tracts such as the SLF, IFOF and in particular the UF show later peak FA values suggesting more delayed developmental maturation (Lebel et al., 2012; Sawiak et al., 2018; Slater et al., 2019) and we did not observe altered FD within these tracts. It may therefore be possible that FBA metrics within an older sample with conduct problems would identify more pervasive (and severe) microstructural impairments across association white-matter tracts. Likewise to the current study, Grazioplene et al. (2020) found no effect of childhood aggression on FD within the UF. However, we did identify that conduct problems were associated with reduced FC within the UF, suggesting that the macroscopic cross-section of fibres may be a more sensitive index for detecting developmental alterations to white matter microstructure within the UF.

A strength of the current study is the relatively narrow age range at each of the two imaging time-points, which allowed for a more focused analysis of the effect of conduct problems and white-matter microstructural properties within development. We also examined white-matter microstructural development across these two time-points, and while conduct problems were related to white-matter alterations at each time-point, we observed no effects of change in conduct difficulties on longitudinal white-matter development between time-points. This was contrary to expectation but may reflect the limited duration between imaging time-points (approximately 16 months) and a longer gap may have allowed for greater developmental differences to emerge in relation to conduct difficulties as the majority of the sample showed no change (42.4%) or a single point change (decrease, 16.8%; increase, 17.6%) from their initial conduct problems score. A further strength of the current study is that we employed a dimensional approach that allowed us to explore the specificity of our findings to conduct problems; this approach is consistent with contemporary conceptualisations of antisocial behaviour as a heterogeneous construct that varies in severity and encapsulates a wide range of externalising and internalising difficulties (Raine, 2018). Importantly, we identified that reduced FD across the fornix and the ILF was primarily driven by increased conduct difficulties rather than emotional, hyperactivity, peer or prosocial problems. This finding is consistent with Grazioplene et al. (2020) who reported that childhood aggression was linked with lower FD and neither anxiety nor callous-unemotional symptoms – the prosocial scale of the SDQ, as used in the current study, has been previously reverse scored to index callous-unemotional traits (Dadds, Fraser, Frost, & Hawes, 2005; Viding, Blair, Moffitt, & Plomin, 2005) – affected this relationship. However, we also found that reduced FC within the UF was not uniquely predicted by either conduct, emotional, hyperactivity, peer or prosocial problems; this may suggest that shared variance across these neurodevelopmental dimensions potentially explains altered FC in development.

Overall, we found that childhood conduct difficulties were related to reductions in white matter fibre density and morphology within the fornix, ILF and UF. Our findings suggest that the development of specific white matter fibre pathways underpinning socio-emotional functioning are related to early conduct problems in childhood.

## Acknowledgements

The Children’s Attention Project was funded by the National Medical Health and Research Council of Australia (NHMRC; project grants #1008522 and #1065895). The data was collected within the Developmental Imaging research group, Murdoch Children’s Research Institute and the Children’s MRI Centre, The Royal Children’s Hospital. It was supported by the Murdoch Children’s Research Institute, The Royal Children’s Hospital, The Royal Children’s Hospital Foundation, Department of Paediatrics at The University of Melbourne and the Victorian Government’s Operational Infrastructure Support Program. We thank the RCH Medical Imaging staff for their assistance and expertise in the collection of the MRI data, and we thank all of the children and families for their participation.

